# Molecular dynamics simulation-based investigation of LexA-DNA interaction: affinity-dependent conformational changes and implications for DNA looping

**DOI:** 10.1101/2023.09.18.558249

**Authors:** Anne Schultze, Mehmet Ali Öztürk

## Abstract

**Context:** The binding of proteins to the DNA plays a crucial role in various cellular processes including gene expression. LexA is a bacterial transcription factor that binds to a specific DNA motif, leading to the regulation of numerous genes involved in the DNA damage response. Understanding the structural dynamics and mechanisms of DNA binding by LexA can provide valuable insights into its function and regulatory capabilities. Here, molecular dynamics (MD) simulations are used to investigate how the sequence of the DNA binding motif influences the conformational changes and bending behavior of DNA upon LexA binding. Simulation trajectories reveal that the DNA fragment containing a higher affinity LexA binding motif exhibits more pronounced bending and structural deformations compared to that containing a lower affinity binding motif. Additionally, bending of the protein itself is also observed in the presence of the DNA fragment with the higher affinity LexA binding motif. Our results confirm previous reports of LexA-induced DNA bending and shed further light on the structural details of the DNA bending mechanisms.

**Methods:** MD simulations are employed to investigate the interaction between LexA and 70 bp-long DNA fragments that differ in the LexA binding motif. The GROMACS simulation package with the AMBER14SB force field for proteins and the AMBERparmbsc1 force field for nucleic acids is used to conduct 100 ns-long all-atom simulations. The solvent is set up with the TIP3P water model and the simulation box is filled with ions to reach 0.15 M salt concentration.

## 1 Introduction

The process of transcription includes an interplay of various components, which makes it a challenging topic to study in detail. Due to the capability of transcription factors (TFs) to specifically recognize and bind to DNA sequences, they are considered as the main regulators of transcription. TFs can bind to a subset of DNA sequences with the strength of the binding depending on the matching of the presented DNA sequence. Thus, greater matching of the DNA motif leads to a higher binding affinity [1]. Regarding the event of transcription itself, DNA has an impact on its efficiency, too. DNA looping effectiveness is a key factor for gene expression levels and is tightly linked to DNA flexibility [2]. The looping can bring TFs from distal binding sites into close proximity of other TFs bound to the DNA, enabling a high concentration of TFs in a specific location [3]. Depending on the positioning of the gathered proteins to the transcriptional starting site, the looping can induce repression or activation of transcription[4], [5]. So far it is known that the DNAs’ length as well as the bendability and flexibility of the DNA sequence itself do have an influence on the looping efficiency [6]. Studies have been conducted regarding the bending of DNA when DNA binding proteins interact with DNA [7], [8]. For several proteins, like the TATA-box binding protein [9] or the CAP protein [10] it is known that bending of the DNA is induced through protein binding and therefore looping gets enhanced. The amount of looping induced by the binding of a protein can be affected by the binding affinity, too [11].

LexA is a naturally occurring bacterial repressor playing an essential role in the induction of the SOS response occurring in bacteria [12]. The protein binds as a dimer to its responsive elements on the DNA through a winged helix-turn-helix motif [13]. The operator sequence targeted by LexA possesses two palindromic sequences, containing 16 bps, which are bound by the two monomers in LexA. The consensus sequence found for LexA responsive elements is CTGT-N8-ACAG [14] with the length of N8 between the two palindromes being fixed [15]. LexA binds to its operator sequence with different binding affinities. A lot of these sequences and their associated binding affinities have been previously determined [16]. Due to its DNA binding capability, LexA’s DNA binding domain can be used as a synthetic transcriptional activator, when a transactivation domain is fused to it. Consequently, LexA no longer acts as a repressor like it does in the SOS response, but can be used as an activator for tran-scription [17]. In this way, gene expression in synthetic biology can be modified in more detail to obtain desired output. For this purpose, the activation domains of several proteins have previously been fused to it (for example GAL4 [18] or VP16 [19]) [20].

It was previously shown in crystal structures that a slight bending of short flanking DNA regions can happen through interaction with LexA, when tight binding is happening [21]. However, no overall investigation was taken out regarding the affinity dependency of bending. Neither were long flanking regions included in the research before.

In order to investigate how changes in binding affinity of the LexA-DNA interaction influence the conformational changes of LexA and DNA, we employed MD simulations on the LexA homodimer - DNA macromolecular complex. For further mechanistic understanding of the effects of LexA in the DNA looping events occurring during transcription, we conducted analyses including RMSD, RMSF and hydrogen bond calculations. Furthermore, the dynamics of LexA and the DNA are further investigated by using distance analysis and calculations on DNA parameters, like the major and minor grooves as well as the roll, tilt and twist angles.

## 2 Methods

### 2.1 Preparation of LexA and DNA structures and docking of the structures

Four different systems of LexA (PDB ID: 3JSP) [22] in complex with two different DNA fragments in B-form and these respective DNA fragments in an unbound state are studied using all-atom MD simulations. The definition of systems can be drawn from Table 1.

**Table 1:** Systems simulated using MD simulations are listed. The name of the system that is used throughout the descriptions in this paper is given.

| system | K<sub>D</sub> value of DNA binding sequence | name |
| --- | --- | --- |
| DNA only | 0.8 nM | DNA08 |
| LexA bound to DNA | 0.8 nM | complex08 |
| DNA only | 5.64 nM | DNA564 |
| LexA bound to DNA | 5.64 nM | complex564 |

The different DNA fragments consisting of 70 bps are generated using the w3DNA web server [23]. They only differ from each other regarding their binding motif and within this their binding affinity and dissociation constant (K_D_). For the protein-DNA complexes, a complete PDB file of the LexA protein is needed, but since there is no complete crystallized structure of LexA available that includes the flexible linker regions entirely, models obtained through the webserver Swissmodel [24] are used, whereas the protein structure of PDB ID: 3JSP is used as template [21]. To generate the macromolecular complex of the modeled LexA homodimer with the DNA, the 16-mer binding sites of the two respective DNA fragments are modeled separately with w3DNA and then docked to the protein structure using the HDOCK web server [25]. By using Pymol [26], the short sequence is replaced by the 70-mer DNA manually. To make the simulation package recognize the DNA properly the first phosphate atoms are deleted from the PDB files.

### 2.2 MD simulation protocol

For the MD simulations, access is granted to the bwForCluster BinAC, offering 236 compute nodes, 62 GPU nodes plus several special purpose nodes for login, interactive jobs, etc. (https://www.binac.unituebingen.de/). All MD simulations are performed with the GROMACS simulation package, version 2022.4. [27]. The AMBER14SB [28] force field is used for the calculation of protein parameters and the force field for nucleic acids is the AMBERparmbsc1 [29]. The systems are explicitly solved using dodecahedron boxes, filled with TIP3P water. The cell borders are set to at least 10 angstroms from the nearest protein atom. Thus, the boxes reach a size around 26 x 26 x 26 nm containing 1205919 (only DNA) to 1206196 (complex) atoms. Periodic boundary conditions are applied. The systems are first neutralized and with added Na+ (1254 ions) / Cl- (1111 ions) counter ions a physiological salt concentration of 0.15 M is reached. Energy minimization using the steepest descent algorithm for a maximum number of 50000 steps to perform and a step size of 0.01 ps is proceeded. In the following, 100 ps of NVT equilibration runs are done to raise the temperature to 300 K, while the volume is kept constant. This step is followed by 100 ps of NPT equilibration where temperature (300 K) and pressure (1 bar) are set constant. Temperature is controlled by a modified Berendsen thermostat [30] and a coupling constant of 0.1 ps and the pressure is controlled by an isotropic Parrinello-Rahman barostat [31] with a coupling constant of 2 ps. All bonds involving hydrogen atoms are constrained with the LINCS algorithm [32] allowing to set the time step to 2 fs. Electrostatic interactions are treated with the Particle Mesh Ewald [33] summation method using the Verlet cutoff scheme [34] with a short-range electrostatic cutoff of 1.0 nm and a short-range van der Waals cutoff of 1.0 nm. The production runs are 100 ns long.

### 2.3 Trajectory analysis

To analyze the trajectories the solvent is removed from the trajectory files to reduce the file size and make handling of the trajectories easier. To visualize the output, the Visual Molecular Dynamics (VMD) [35] tool is used. The structural information obtained during the MD simulation is analyzed in terms of root mean square deviation (RMSD), root mean square fluctuation (RMSF) and distance analysis with the MDtraj python package [36]. The number of distinct hydrogen bonds at the protein-DNA interface is analyzed with the built-in analysis tools in the GROMACS package. The DNA parameters are calculated using PYTRAJ, which represents the python version of the CPPTRAJ tool [37].

## 3 Results and Discussion

The DNA sequences that are docked to LexA prior to the MD simulations differ in their LexA binding sequence (Figure 1.a). The DNA with a K_D_ value of 0.8 nM is bound more tightly to LexA compared to the DNA with a K_D_ value of 5.64 nM [21] .

**Fig. 1:**
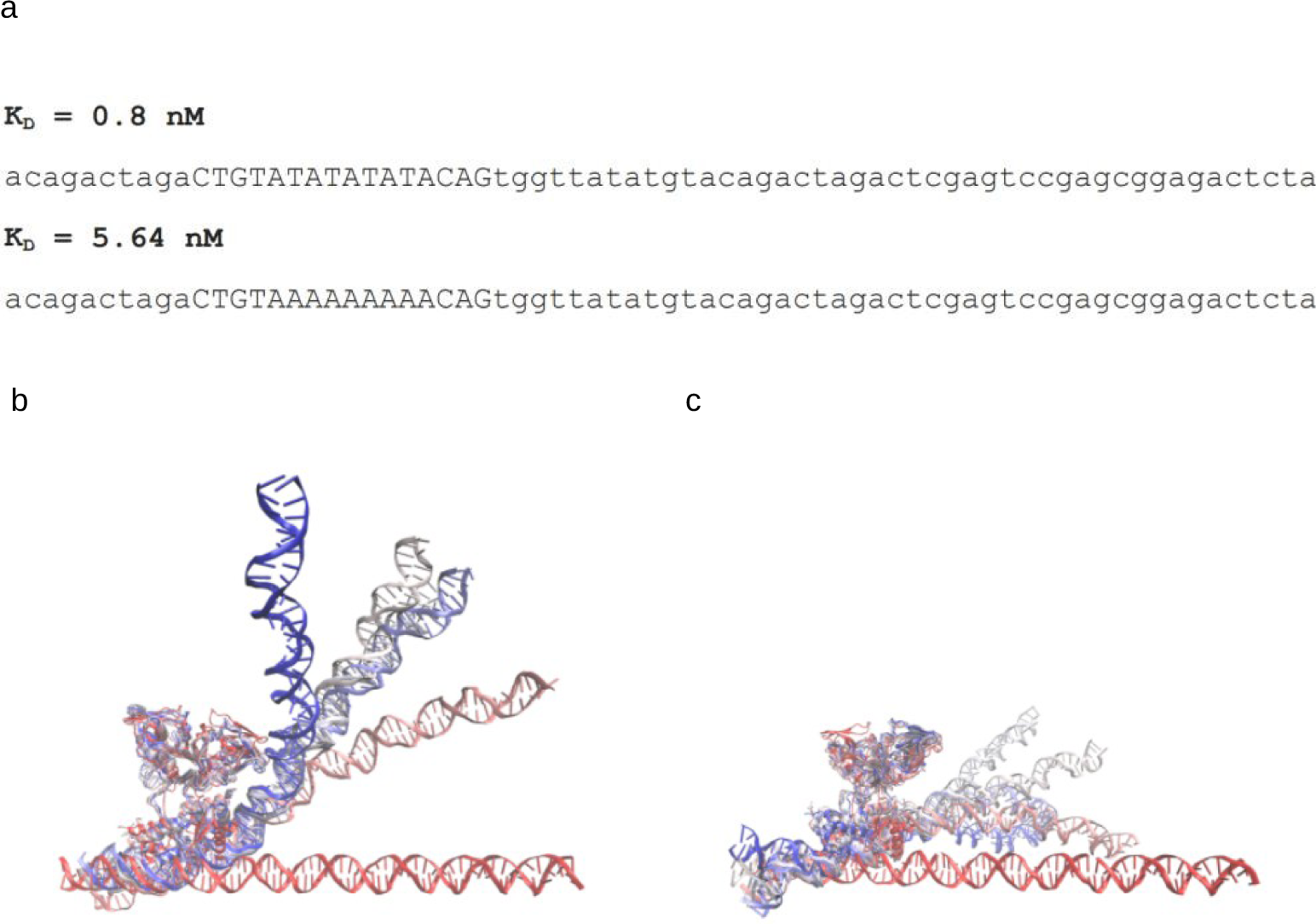
Analyzed DNA sequences with different binding motifs and the visual analysis of the LexA homodimer-DNA simulation runs a) The nucleotide sequences of the DNA fragments used as template for the analyzed systems are given. Bases written in capital letters show the LexA binding sequence which differs between the two DNA fragments. Bases written in small letters show the flanking sites of the analyzed DNA sequences which are similar in both fragments. The K_D_ values describe the dissociation constants of the binding motifs in the respective DNA sequences. b and c) The movement of the systems where the DNA is bound by LexA is visualized using snapshots of every 20 ns frame of the trajectories with the aligned structure of the protein. The input structure (first frame of the simulation) is shown in red, which turns to blue (final conformation at the end of each trajectory). Snapshots during the simulation show gradually changing colors from red to blue. b) 100ns trajectory of complex08. c) 100ns trajectory of complex564

To explore the potential correlation of the affinity between LexA and its operator sequence and DNA bending, MD simulations and several downstream analyses were performed. For better comparison of the moving behavior the two different DNA fragments were simulated in a bound and in an unbound state, which leads in total to four different systems to be investigated (see Table 1). To give an overview over the trajectories Figure 1.b and c depict the structural movement of the LexA-DNA complexes over the simulated timespan. The DNA in complex08 bends strongly in the direction of LexA with its long flanking sequence (Figure 1.b). Similar movement cannot be detected regarding the simulation of complex564 (Figure 1.c). To further analyze the behavior of LexA and the DNA, quantitative analyses are done in the following.

### 3.1 MD simulations reveal structural differences of the LexA – DNA complexes dependent on presented DNA motifs

To analyze the overall behavior of the different systems an RMSD (root mean square deviation) analysis is conducted. This analysis gives an insight on how much conformational convergence exists in the simulation, how steady the states are throughout the simulation and it further visualizes the differences in structural orientation, moving and stabilization. Within the scope of the RMSD analysis a notable distinction between the two formed complexes can be shown (Figure 2.a). The stability of the bound protein depends on the binding motif of the DNA. LexA bound to the high affinity binding motif shows lower RMSD values and stabilizes quickly after the simulation starts. LexA bound to the low affinity binding site does not stabilize during the entire simulated timespan and reaches values double as high as those calculated for the other complex.

**Fig. 2:**
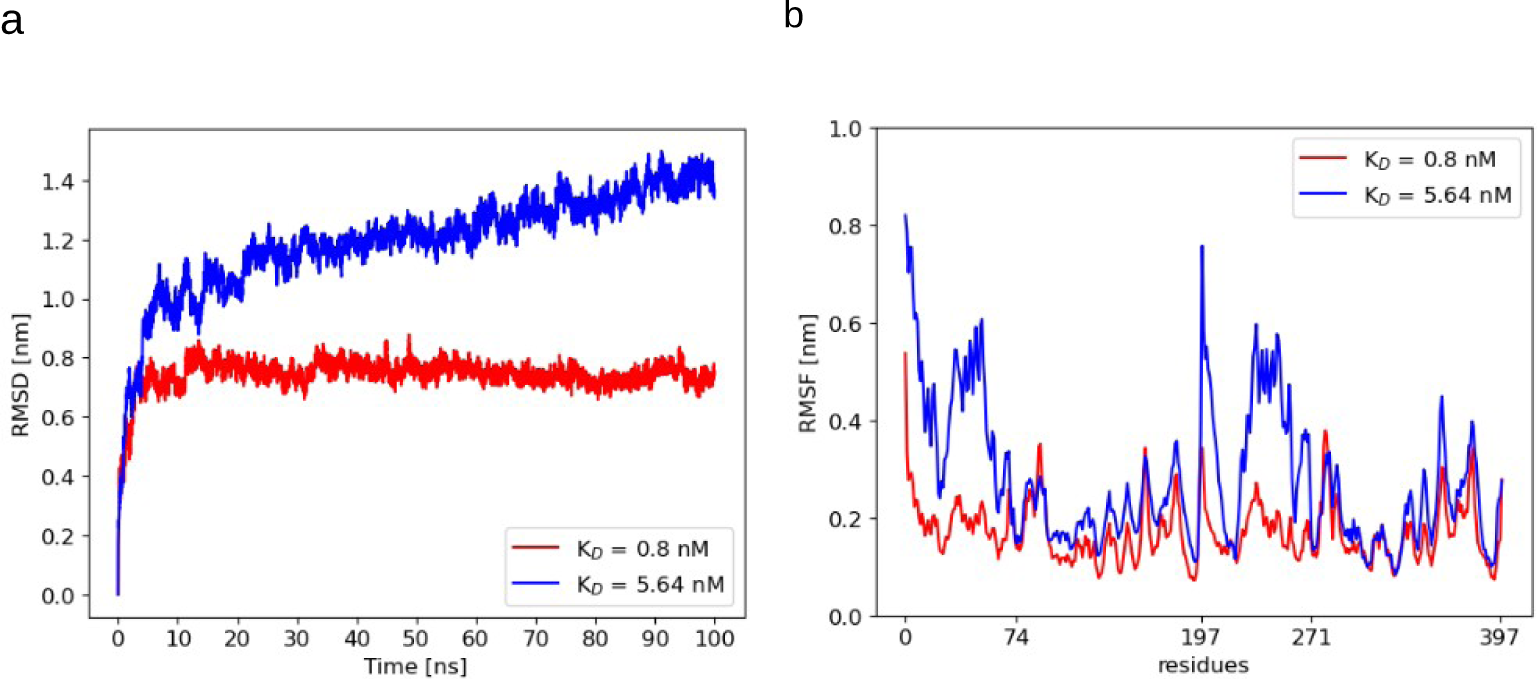
RMSD and RMSF of LexA bound to the DNA a) The RMSD of the protein LexA is shown for both trajectories including LexA: complex08 (red) and complex564 (blue). The frames considered in the calculations are frames taken of every nanosecond of the respective trajectories of 100 ns each. b) The RMSF values of LexA in complex08 (red) and complex564 (blue) are plotted. The DNA-binding domain includes residue 0 to 74 (first monomer) and residue 197 to 271 (second monomer). Residues 75 to 196 (first monomer) and 272 to 397 (second monomer) are the residues of the C-terminal domain.

To go more into detail on the flexibility of single residues of LexA, the RMSF (root mean square fluctuations) are calculated to analyze the residue specific flexibility of the systems, focusing on flexible regions (Figure 2.b). The RMSF analysis supports the theory that binding to the high affinity binding site stabilizes the formed complex more than the low affinity binding site, as the overall calculated fluctuation values for the protein as well as for the DNA sequence from the complex formed with the low affinity binding motif are higher. The RMSF of the protein residues depict the differences regarding the flexibility of the residues in the two domains incorporated by LexA. The DNA binding domain of the protein (residue 0 to 74 and residue 197 to 271) connected to the low affinity binding site shows elevated values compared to the protein from the other complex which depicts the impact of strong binding on the flexibility of the residues.

The RMSF of LexA is followed by the same analysis of the DNA fragments to show flexible regions and differences in behavior of the nucleotide sequences (Figure 3). It depicts that the DNA sequence possessing the high affinity binding motif is more flexible in an unbound state (Figure 3.b). When protein binding occurs, this switches and the DNA with the low affinity binding site shows an overall greater flexibility, except regarding the nucleotides of the binding motif. In addition, the RMSF shows that the flanking regions of the DNA sequences are the more flexible ones and that the DNA sequences show a pattern regarding the position of flexible residues over the entire sequence. Considering the sequences that are used for this analysis no pattern regarding A-T or G-C-rich regions can explain the fluctuation pattern (Figure 3.a). The TA-rich regions (present in the binding motif of complex08) which are known to be very flexible [38] also show a great flexibility in the simulations, whereas the AA-rich regions (included in the binding motif of complex564) show less flexibility.

**Fig. 3:**
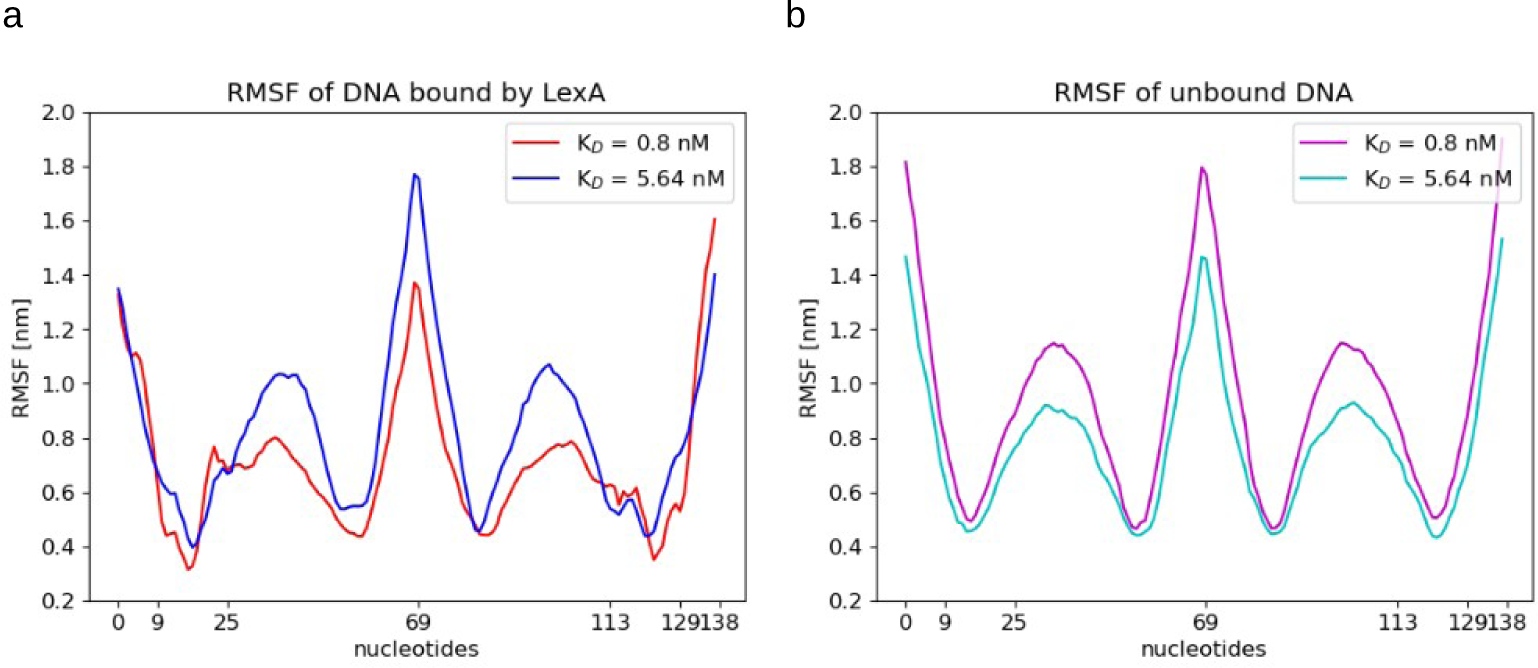
RMSF of the DNA The first strand consists of the residues 0 to 69, the second one of the residues 70 to 138. Residue 9 to 25 and 113 to 129 show the parts of the DNA where the LexA binding motif appears. a) bound DNA in complex08 (red) and complex564 (blue). b) unbound DNA08 (magenta) and DNA564 (cyan)

The hydrogen bond analysis strengthens the existing knowledge about high and low affinity binding sites and the interactions appearing between the DNA and the protein when binding occurs [39]. The complex formed with the DNA presenting the high affinity binding site shows more formed bonds during the entire simulation timespan (see Supplementary Figure A1).

It is well known that the binding motif and the affinity of the protein binding to it, have a huge impact on the stability of the complex and therefore also on the gene expression levels [40]. The results of our analysis are consistent with this knowledge. Changing single bps alters the binding affinity and therefore the whole interaction of the protein with the complex. The TA-rich regions which are known to be very flexible also show a great flexibility in the simulations [38], whereas the AA-rich regions show less flexibility. Due to a reduction of the enthalpic cost for bending because of the looser stacking of TA-sequences, higher flexibility is reached in these regions [41]. This prior knowledge is consistent with our results.

### 3.2 Bending of the DNA is induced through LexA binding

To analyze the bending of the 70 base pair-long DNA fragments, distances on the DNA that approximate the sizes of major and minor grooves are defined [42]. They are specified by choosing the phosphate atoms of every tenth base pair on the backbone of the DNA, which leads to seven major grooves and six minor grooves (Figure 4.a).

**Fig. 4:**
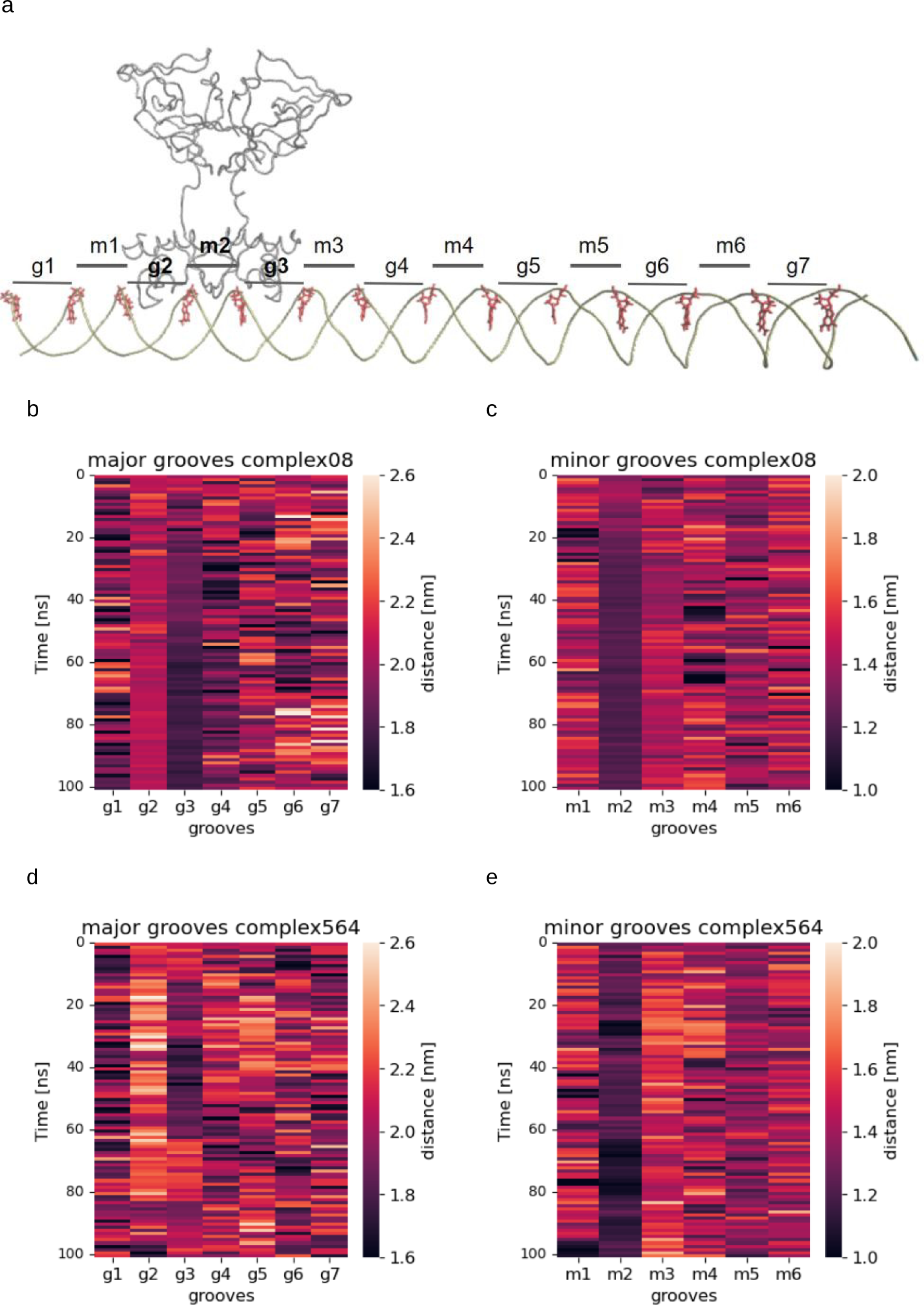
Residues for the definition of grooves and major and minor groove distances of DNA fragments bound by LexA a) The position of the defined residues for the groove calculations is shown as well as the defined groove numbers. Minor grooves are named m1 to m6, whereas major grooves are called g1 to g7. b) - e) The distances calculated between the phosphate atoms of every 10 base pairs on the 70 bp long DNA fragments to define major and minor grooves on the DNA are depicted over time. Frames of every nanosecond over the 100 ns simulation are considered. b) major groove distances of the DNA in complex08. c) minor groove distances of the DNA in complex08. d) major groove distances of the DNA in complex564 e) minor groove distances of the DNA in complex564

The groove analysis shows a striking difference between the two complexes (Figure 4.b-e). It displays that in the protein binding region bending occurs and is dependent on whether LexA binds with high or low affinity to the DNA. The high affinity binding motif leads to stronger bending of the DNA. The major groove comparison shows huge dissimilarities between the two complexes. The major grooves g2 and g3 and the minor groove m3 are narrower in complex08 than in complex564. The narrow grooves indicate bending of the DNA in this region in the complex formed of LexA with the DNA incorporating the high affinity binding motif. Moreover, an asymmetric bending of the DNA is visible in complex08, as major groove g3 becomes narrower than major groove g2, as well as minor groove m3 is stabilized more than minor groove m1. This shows an extent of the bending occurring on the long site of the flanking DNA sequence that is identical in all systems. The strong bending of minor groove m2 in both complexes is suggested to make a binding of the wingtips present in the LexA dimer possible across the minor groove [21]. These specific patterns are not visible for the unbound DNA fragments (see Supplementary Figure A2).

To analyze the bending of the DNA in more detail, the roll, tilt and twist angles are calculated between all 70 bps. These are parameters defined for the description of the behavior of DNA [43]. The roll angle describes bending by showing the rotation between two neighboring bps along the z-axis. The tilt angle shows bending along the y-axis and the twist angle defines bending happening along the x-axis, which already occurs through the natural helical structure, but can be influenced by bending, too [44].

The roll angle calculations show bending in the site of the operator sequence (Figure 5.a,b). The angles calculated for the complex with the DNA presenting the high affinity binding motif are bigger than those from the complex with the low affinity binding motif. This is consistent with the fact that strong binding leads to more conformational change in the DNA [45]. No such pattern is visible regarding the unbound DNA fragments (see Supplementary Figure A3.a,b). The twist angle calculations indicate bending in both complexes, but stronger twisting occurs regarding the complex formed with the DNA presenting the high affinity binding motif (Figure 5.c,d). The twisting occurs primarily on the site of the operator sequence adjacent to the long flanking DNA sequence (bp 18 to 26), which aligns with the results from the groove size analysis. Again, this pattern is only visible in the fragments bound to LexA, not in the unbound fragments (see Supplementary Figure A3.c,d). The tilt angle calculation does not show any differences regarding the behavior of the bound and unbound DNA fragments (see Supplementary Figure A4).

**Fig. 5:**
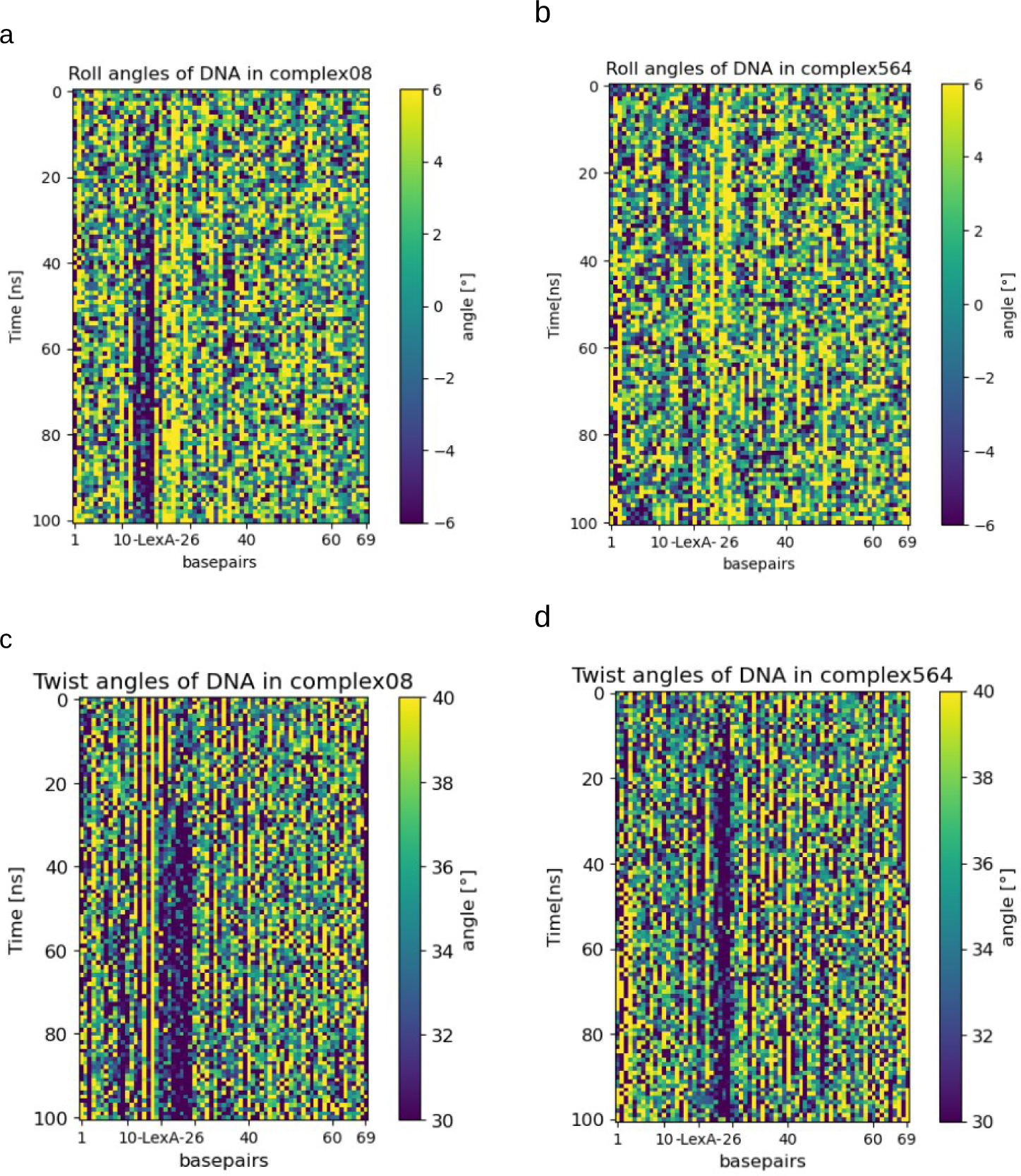
Roll and twist angles of the LexA bound DNA fragments The roll and the twist angles calculated between the single bps of the DNA sequences for the analyzed systems of bound DNA fragments are depicted over time. Frames of every nanosecond over the 100 ns timespan are considered. a) Roll angles of DNA in complex08. b) Roll angles of DNA in complex564. c) Twist angles of DNA in complex08. d) Twist angles of DNA in complex564. To better visualize the direction of bending in the heatmaps, the scales are adjusted to lower values, as the angles can span a large range. This means, a value of 20, for example, is represented with the same color as a value of 6, while a value of -20 is depicted with the same color as a value of -6

### 3.3 Asymmetric movement of LexA dimer occurs through binding to high affinity DNA motifs

To visualize the movement within LexA when bound to DNA, distance measurements between the C-alpha atoms of residues at the outer ends of the protein in both - the DNA binding domain and the C-terminal domain - are performed and plotted over time (Figure 6).

**Fig. 6:**
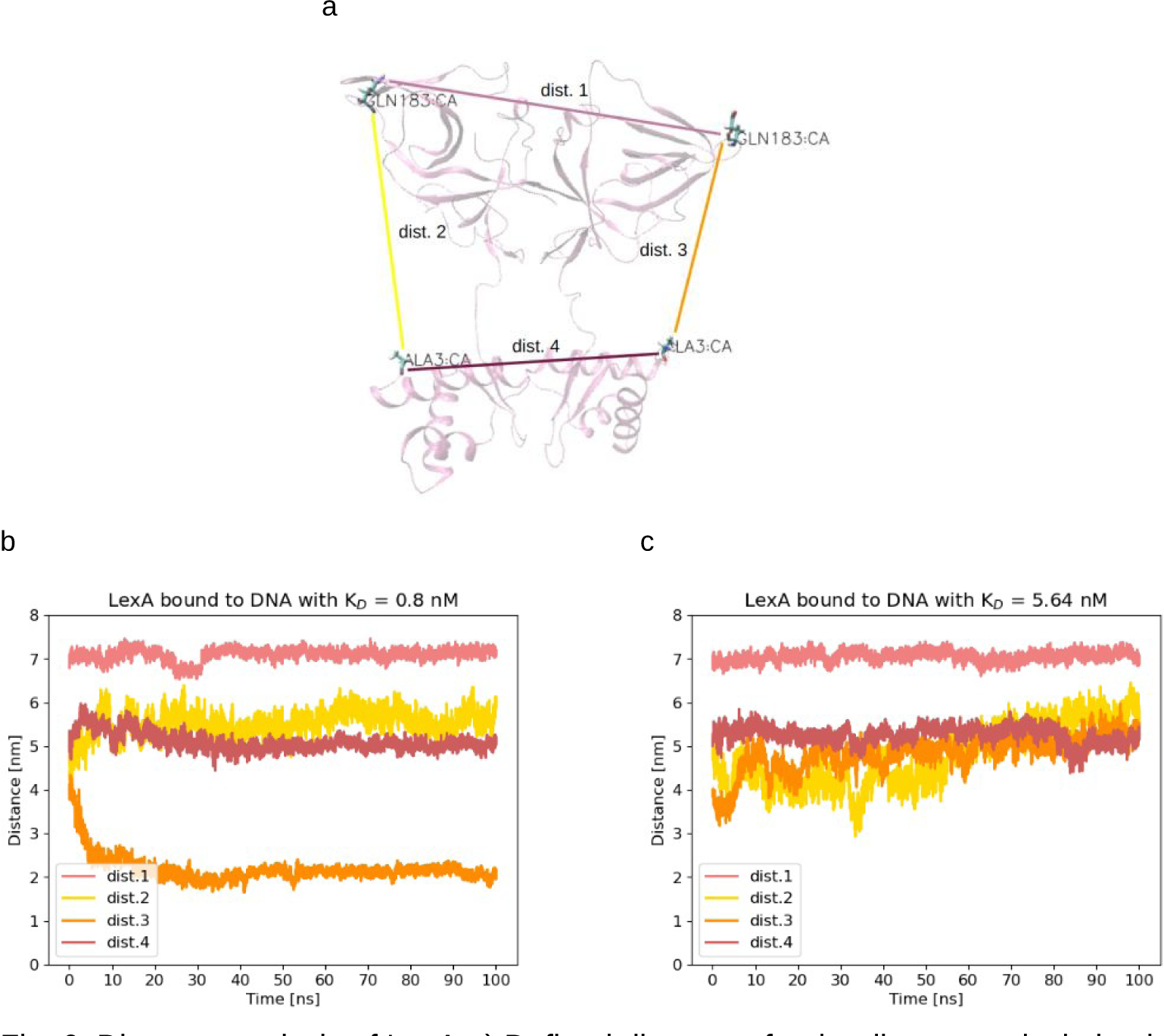
Distance analysis of LexA a) Defined distances for the distance calculation in LexA: Dist.1 describes the distance between the carbon atoms of two glutamine residues of each monomer in the C-terminal domain of LexA. Dist.2 describes the distance between the glutamine and the aspartic acid in the first monomer. Dist.3 shows the same distance in the second monomer. Dist.4 shows the distance between two aspartic acid C-alpha atoms in the DNA-binding domain of each monomer. b) the distances in LexA in the simulation of complex08 are shown. c) the distances of LexA in complex564 are shown.

The distance analysis of the protein reveals a different behavior of LexA when bound to the high affinity binding motif compared with the low affinity binding motif (Figure 6.b,c). The distances between the two monomers (dist.1 and dist.4) do not change much over the simulated time span, which supports previous findings on the dimer stabilized through the binding to DNA [46]. Through the flexible linker between the N and C-terminal domains of the protein, these can easily rotate, even when bound to DNA. Thus dist.1 and dist.3 are the most changing ones. The protein in complex with a low affinity binding site shows rather a rotation of its two domains and no bending, as the values of dist.2 and dist.3 both rise during the simulated timespan (Figure 6.c). This movement of LexA when binding to the DNA has been previously discussed [47].

As dist.3 in Figure 6.b indicates through dropping values, LexA bound to the high affinity motif bends towards the long site of the DNA. Especially residues 162, 163 and 182 and 183 are in closest proximity to the DNA. This could suggest that the complex is not only stabilized at the binding site, but actually another site of LexA interacts with the DNA for further stabilizing the complex. This could indicate a second DNA binding site within LexA which gets contacted only after the first DNA motif is tightly bound.

## 4 Conclusion

By using MD simulations we could show that LexA is indeed inducing bending of the DNA. Furthermore, we demonstrate that this induced behavior depends strongly on the strength of binding. Binding of a high affinity binding motif leads to strong bending, whereas the binding of a low affinity binding site does not show such a behavior.

In prior research, it was shown that the binding of LexA induces bending of the DNA in the operator region when bound to the caa operator, but less when bound to the reca operator. These assays were performed in 1988 with gel electrophoresis shift assays [48]. Nowadays, these experiments are not done anymore to prove DNA bending, as they only provide poorly reliable results. The simulations done as part of this work, are not done with the same operator sequences that were used in 1988, but do indicate different bending behavior depending on the motif LexA binds, too. More recent work on LexA-induced DNA bending was done using X-ray crystallography [22]. The bending of the DNA within the region of the consensus sequence can be confirmed by the results obtained from the MD simulations. Nevertheless, the structures analyzed by Zhang et al. do not contain DNA fractions as long as the ones used for the simulations done here. The previous investigations only show bending in the region directly interacting with LexA but not regarding the shape of the flanking regions. The results of the MD simulations show that the positioning and conformation of the entire DNA sequence is influenced by the strength of LexA binding and thus the operator sequence present. Prior research indicated that the flexibility of flanking regions can have an impact on the stabilization of protein-DNA complexes [15], but not that the binding motif has an impact on the flanking regions.

Direct and indirect shape readout are very important types of investigation to further understand the timing of events for transcription. A rising question is if the DNA is prebent and thus enhances protein binding (called conformational capture) or if the binding of the protein itself induces bending [11]. Looking at the analysis of the unbound DNA, it can be said that the bending does not occur without LexA docked to it. This indicates that the bending is induced through protein binding. It is well known that LexA relies heavily on base readout rather than shape readout, while the stabilization of the complex relies on shape readout of the flanking regions of the DNA [49]. These prior findings are supported by the results of the MD simulations presented here.

The obtained results deepen our understanding of the molecular basis of LexA-DNA recognition and provide a foundation for future studies aiming to engineer synthetic transcription factors for defined gene expression levels and application in therapeutic approaches. Moreover this study demonstrates the power of molecular dynamics simulations in elucidating the dynamic aspects of protein-DNA interactions and being able to simulate even relatively longer DNA fragments.

Supplementary information. Additional figures of further analysis can be found in the Supporting Information.

## Acknowledgments

We thank Barbara Di Ventura for critical discussions and providing the funding for this work. We also thank the High Performance and Cloud Computing Group at the Zentrum für Datenverarbeitung of the University of Tübingen, the state of Baden-Württemberg through bwHPC and the German Research Foundation (DFG) through grant no INST 37/935-1 FUGG.

Funding. This work was funded by the Deutsche Forschungsgemeinschaft (DFG) under Germany’s Excellence Strategy through EXC294 (BIOSS—Center for Biological Signalling Studies), and EXC2189 (CIBSS—Centre for Integrative Biological Signalling Studies, Project ID 390939984). M.A.O. additionally received funding from the European Research Council (ERC) under the European Union’s Horizon 2020 research and innovation program (ERC Co grant to Barbara Di Ventura; Grant agreement No. 101002044).

Authors’ contributions. A. S. conducted all simulations, applied trajectory analyses and prepared the manuscript draft. M.A.O. overall supervision of the project, analysis of results, editing the manuscript draft.

Conflict of Interest. The authors declare no conflicts of interest.

## Appendix A Supplementary Information

**Fig. A1:**
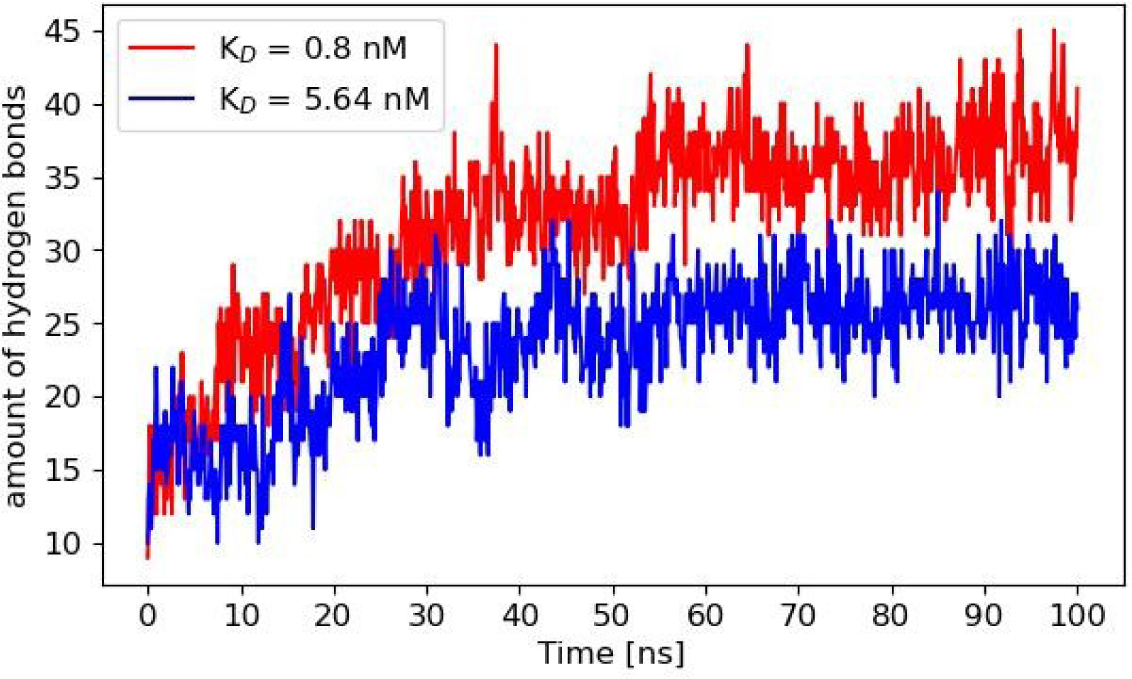
Hydrogen bonds between the DNA and LexA The amount of hydrogen bonds existing between LexA and the DNA is depicted over the simulation time in red for complex08 and blue for complex564

**Fig. A2:**
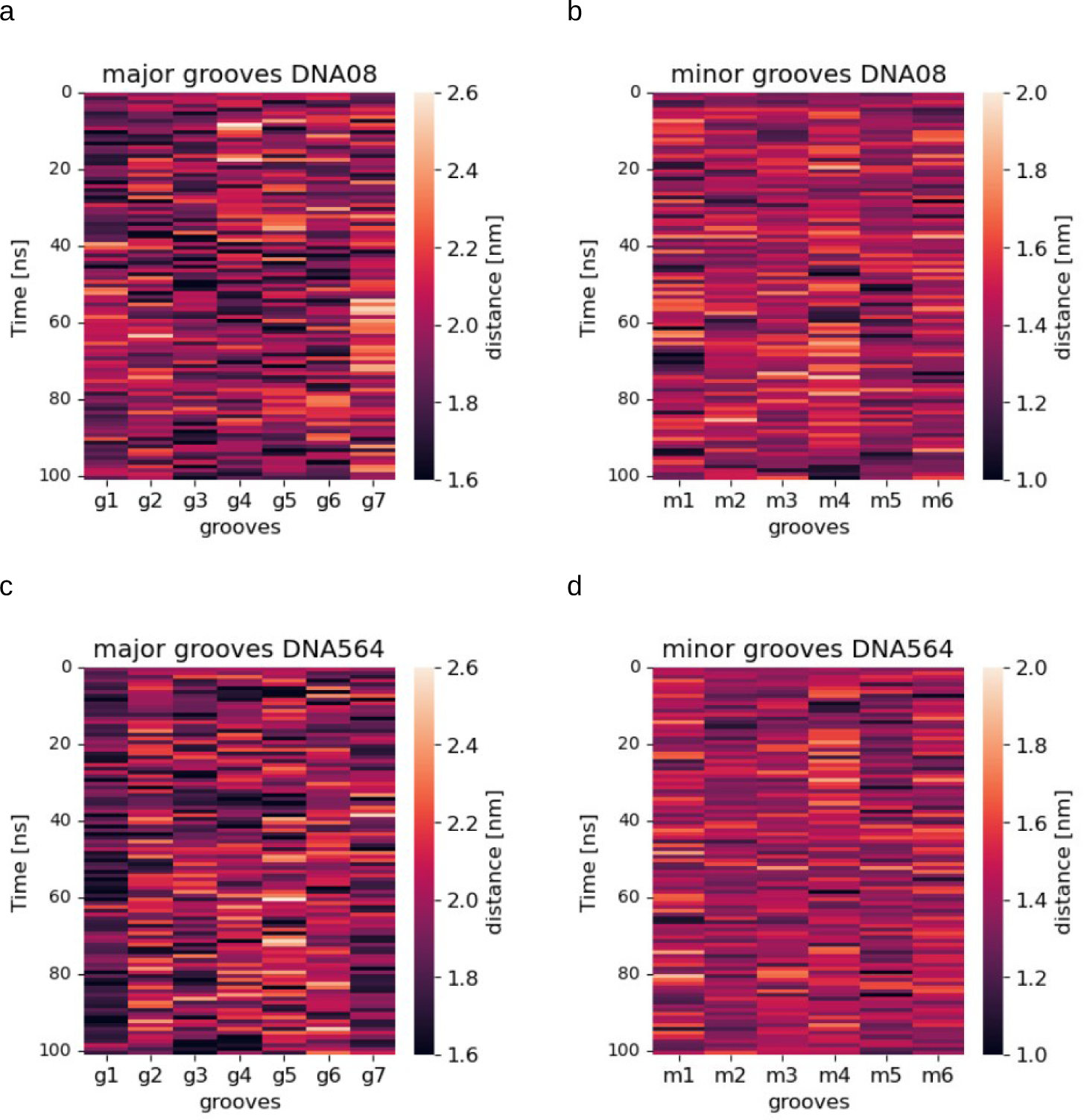
Major and minor groove distances of the unbound DNA fragments The distances calculated between the phosphate atoms of every 10th bp on the 70 bp long DNA fragments to define major and minor grooves on the DNA are depicted over time. a) major groove distances of DNA08 b) minor groove distances of DNA08 c) major groove distances of DNA564 d) minor groove distances of DNA564

**Fig. A3:**
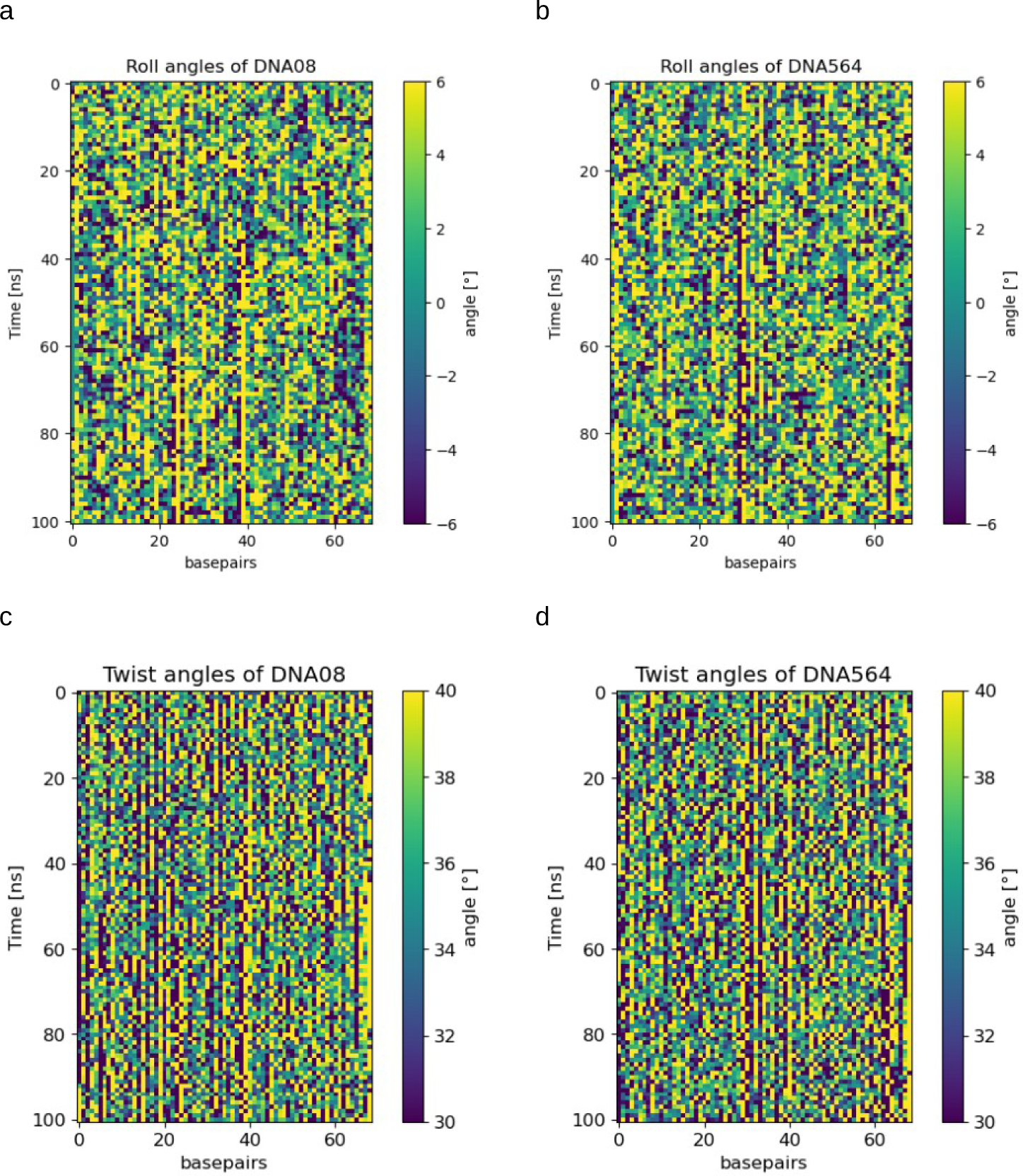
Twist angles of the unbound DNA fragments The roll and twist angles between the bps of the DNA sequences for the analyzed unbound DNA fragments are depicted over time with frmaes considered every 100 ns. a) roll angles of DNA in complex08. b) roll angles of DNA in DNA08. c) Twist angles of DNA in complex564. d) Twist angles of DNA in DNA564

**Fig. A4:**
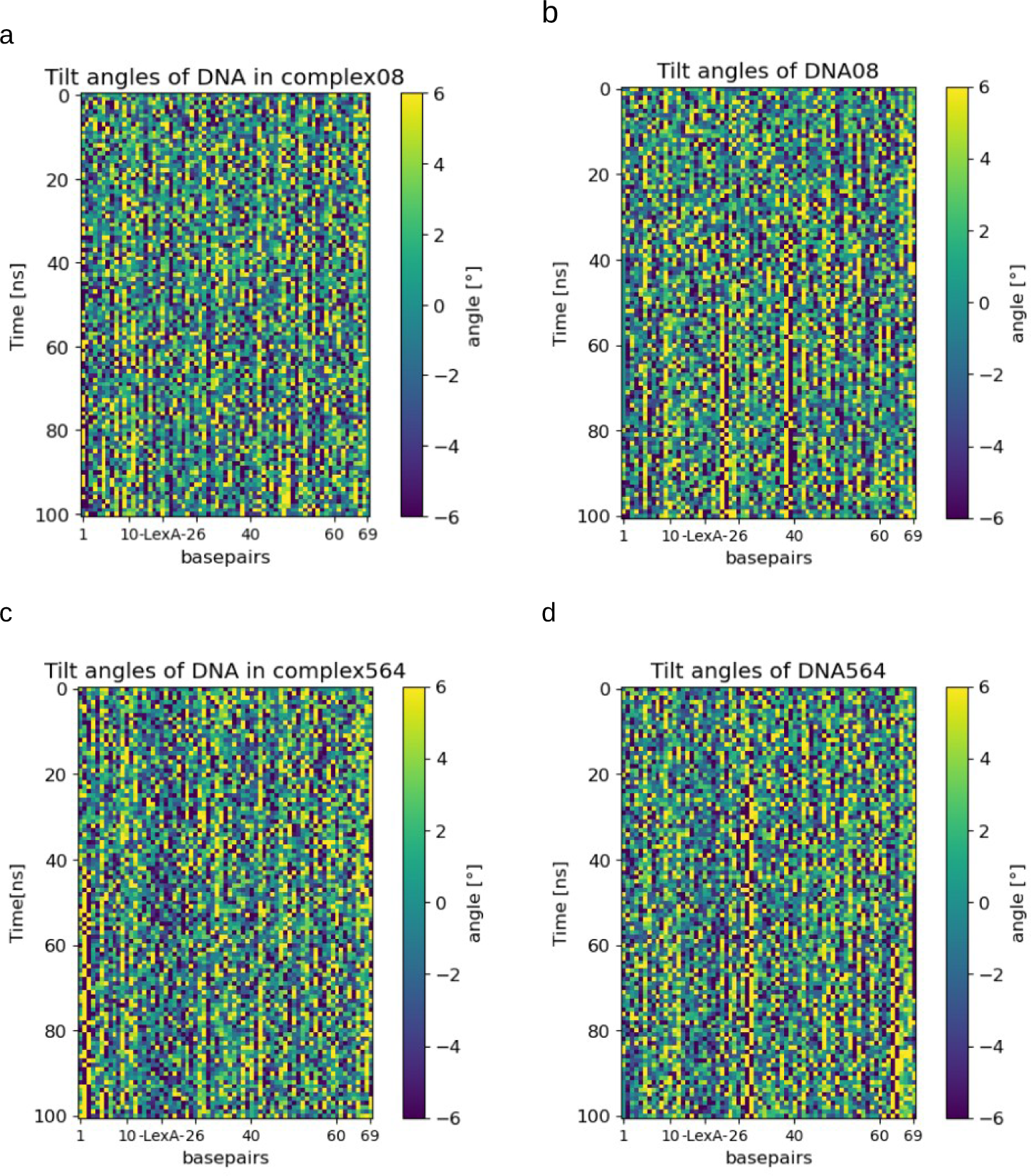
Tilt angles of the DNA The tilt angles between the bps of the DNA sequences for the four analyzed systems are depicted over time. a) DNA in complex08. b) DNA in DNA08. c) DNA in complex564. d) DNA in DNA564

